# Increased contact between lipid droplets and mitochondria in skeletal muscles of male elite endurance athletes

**DOI:** 10.1101/2025.02.05.636640

**Authors:** Joachim Nielsen, Kristine Grøsfjeld Petersen, Martin Eisemann de Almeida, Sam O. Shepherd, Britt Christensen, Maria Houborg Petersen, Kurt Højlund, Niels Ørtenblad, Kasper Degn Gejl

## Abstract

Endurance athletes exhibit higher skeletal muscle mitochondrial and lipid droplet (LD) content compared to recreationally active individuals, along with greater whole-body oxygen uptake and maximal fat oxidation rates. In this study, we investigated if these differences manifest in a greater LD-mitochondria contact and how this may relate to the organelles’ size, shape, and numerical densities. We obtained skeletal muscle biopsies from 17 male elite triathletes and road cyclists and 7 recreationally active men. Using quantitative transmission electron microscopy, we found that the endurance athletes had 2-3-fold greater LD-mitochondria contact length than the recreationally active individuals. This was related to higher numerical densities of both mitochondria and LDs in the intermyofibrillar space. Adding data from untrained individuals with equally high intermyofibrillar LD density as the endurance athletes revealed a 24% greater LD-mitochondria contact length in the endurance athletes. We observed small trivial differences in the shape of both organelles between populations. However, large mitochondrial profiles were more elongated and irregular in shape compared to small mitochondrial profiles, while large LD profiles were more circular and less irregular than small LD profiles. Within the group of athletes, large intermyofibrillar LD profiles correlated with a high fraction of PLIN5-positive LDs and their maximal fat oxidation rate was positively associated with an interaction between the profile size of both intermyofibrillar LDs and mitochondria. In conclusion, male endurance athletes have a greater LD-mitochondria contact than recreationally active and untrained individuals. This muscular phenotype is restricted to the intermyofibrillar space and to fibres rich in mitochondria.

## Introduction

Endurance-trained athletes exhibit a superior fat oxidation capacity and rate at a given exercise intensity compared to sedentary or recreationally active individuals (Klein et al. 1994). This enhancement is achieved through multiple factors, such as an increased transport capacity of fatty acids into the skeletal muscle fibres and further into the matrix of an increased volumetric network of mitochondria (Talanian et al. 2010; Smith et al. 2012), where fatty acids are catabolized via β-oxidation. In addition to the influx of fatty acids from the plasma into muscle fibres, fatty acids can also be mobilized from stored triglycerides within the muscle fibres, a process that is upregulated by endurance training (Alsted et al. 2009; Shepherd et al. 2013). These triglycerides are stored in lipid droplets (LDs) distributed throughout the muscle. It can be reasoned that the physical proximity of LDs to mitochondria influences the rate of fatty acid turnover, thereby providing an additional site of regulation for the whole-body fat oxidation rate and capacity. This rationale is based on the finding that contact between LDs to mitochondria facilitates the delivery of fatty acids to mitochondria (Miner et al. 2023) and that mitochondria associated with LDs exhibit higher respiratory capacity compared to those free of LDs (Kim et al. 2024). The proximity of LDs and mitochondria can be evaluated by membrane-membrane interactions, with transmission electron microscopy of muscle biopsy specimens providing high-resolution quantification of this interaction (de Almeida et al. 2023a).

Most studies have investigated LD-mitochondria interaction binarily by classifying LDs or mitochondrial profiles as either in contact with the other organelle (also termed “touching”) or not. Studies using this approach have shown mixed results regarding a difference between trained and untrained individuals or the presence of a training effect (e.g. Tarnopolsky et al. 2007; Samjoo et al. 2013; de Almeida et al. 2023a), suggesting methodological limitations or a myriad of factors involved. We have recently refined this method by quantifying the length of membrane-to-membrane interaction, and found no differences between untrained lean and obese men with or without type 2 diabetes (de Almeida et al. 2023a). As trained have a higher fat oxidation capacity, this may be associated with a higher LD-mitochondria membrane-to-membrane interaction. However, it remains to be investigated whether highly endurance-trained athletes have a greater length of contact than untrained individuals.

Important factors involved in LD-mitochondria interactivity are subcellular localization (de Almeida et al. 2023a), mitochondrial network characteristics (Bleck et al. 2018), and LD-coating proteins (Shepherd et al. 2012). Specifically, highly trained endurance athletes exhibit an extraordinarily high number of LDs in the intermyofibrillar (IMF) region (i.e., between the myofibrils) (Koh et al. 2017). Furthermore, endurance exercise training and high-intensity interval training have been shown to increase the presence of LDs in this specific region (Devries et al. 2013; de Almeida et al. 2023a). However, the metabolic implications of the increased abundance of LDs in the IMF region in endurance-trained individuals remain uncertain, and it is unknown whether the size, shape and number of LDs influence their membrane-to-membrane interaction with mitochondria.

LDs are coated by several proteins involved in LD turnover and mitochondrial interaction, such as perilipins (PLINs). PLIN5 is abundantly expressed in skeletal muscles, and a growing body of evidence suggests that PLIN5 plays a pivotal role in intracellular lipid homeostasis, including the regulation of lipolysis and lipogenesis (Zhang et al. 2022). Interestingly, PLIN5 has consistently been observed at the interface between LDs and mitochondria, anchoring mitochondria through its C-terminal amino acids (Bosma et al. 2012; Gemmink et al. 2018; Wang et al. 2011). It has been suggested to facilitate the flow of fatty acids to the mitochondria, as the expression of PLIN5 has been shown to correlate with mitochondrial respiration rates of lipid derived substrates (Bosma et al. 2012). Hence, PLIN5 appears to enhance the physical contact between LDs and mitochondria possibly participating in the regulation of fat oxidation during exercise.

The aim of the present study was to investigate whether endurance-trained athletes exhibit a higher contact length between LDs and mitochondria in skeletal muscle fibres than that observed in recreationally active individuals, and furthermore how this would be described by differences in LD and mitochondrial morphology, content, and localization. In addition, within the group of endurance athletes, we examined whether the amount of physical contact between LDs and mitochondria was associated with the content of PLIN5.

## Methods

### Ethical approval

The participants were fully informed of any potential risk associated with the experiments before verbal and written consents were obtained. The Ethics Committee of Southern Denmark approved the study protocols (Project-ID’s: S-20150034 & M-20110035) and the experiments adhered to the standards of the *Declaration of Helsinki*.

### Participants

Seventeen highly trained male triathletes and road cyclists (ET) were enrolled in the study (mean (SD) age 24.1 ± 3.2 years, body weight: 77.1 ± 5.8 kg, V̇O_2max_: 65.1 ± 4.9 ml O_2_ min^-1^ kg^-1^). The 12 male triathletes and 5 road cyclists trained at least 10 hours per week, demonstrated a V̇O_2max_ greater than 60 ml · kg ^-1^ · min^-1^ and had at least 2 years of experience in their discipline. Among the triathletes, six were current members of the Danish national team competing at international Olympic and sprint distances (Olympic Games, World Triathlon Series, World Cups and Continental Cups), 3 participated in national elite competitions (Olympic, ½ ironman and ironman distances), while the remaining 3 competed at a lower level (½ ironman and ironman distances). All five cyclists had A-licenses and competed at the national elite level. In addition, seven healthy, recreationally active men (RA) (age 23.6 ± 2.0 years, body weight: 80.3 ± 9.0 kg, V̇O_2max_: 45.4 ± 5.1 ml O_2_ min^-1^ kg^-1^) were included. The groups of athletes and recreationally active men are sub-groups of individuals from larger projects, from which some data has already been published (Gejl et al. 2017; Nielsen et al. 2017). In Fig. 8, observations from an additional 43 untrained lean men and obese men, with or without type 2 diabetes (age 55.0 ± 6.4 years, body weight: 94.5 ± 14.7 kg, V̇O_2max_: 32.2 ± 7.6 ml O_2_ min^-1^ kg^-1^), were included (Petersen et al. 2022; de Almeida et al. 2023a).

### Muscle biopsies

A muscle biopsy of 100-150 mg was obtained at rest from the *m. vastus lateralis* portion of *m. quadriceps femoris* using 5 mm Bergström needles. A 1 cm incision was made in the middle region of the m. vastus lateralis under local anaesthesia before the biopsy was obtained by the percutaneous needle biopsy technique. Muscle tissue was placed on filter paper upon an ice-cooled Petri dish, blotted and dissected free from fat and connective tissue. The participants refrained from exercise for at least 36 hrs. preceding the biopsy extraction. For the purposes of the present study, one part was used for TEM imaging (ET and RA), one part for confocal microscopy (only ET), and one part was homogenized and used for measurement of enzyme activity and fibre type distribution (only ET).

### V̇O_2_ measurements

Using an electronically braked ergometer (Schoberer Rad Messtechnik (SRM), 117 GmbH Julich, Germany) the ET group conducted a submaximal incremental test followed by a V̇O_2max_ test. The submaximal incremental test was composed of 4-min intervals at an initial workload of 135 W. The workload was elevated by 35 W every fourth minute until the RER value remained above 1.00 for one whole minute. The V̇O_2max_ test was conducted 15 mins after termination of the submaximal test and the initial 2-min workload corresponded to the workload during the penultimate step of the submaximal test. Thereafter the workload was increased by 25 W every minute until exhaustion. The ergometer was calibrated before each test. In the RA group, V̇O_2max_ was determined using a Monark ergometer bicycle (Monark Ergomedic 828E; Monark, Varberg, Sweden). This test was performed at a constant pedaling rate of 70 rpm, starting at 140 watts. Hereafter, the workload was increased with 35 watt every minute until exhaustion.

In both the ET and RA participants, V̇O2 and V̇CO2 were continuously sampled every 10 seconds using an Oxygraf CPET OEM system (AMIS 2001, Innovision, Glamsbjerg, Denmark), based on pulmonary ventilation and expiratory CO2 and O2 concentrations. Prior to each test, the gas analyzer was calibrated with two standard mixtures of gases containing 21 and 15% of O_2_ and 0 and 5% of CO_2_. Ventilation sensors were calibrated manually with a 3L syringe. The highest mean 30-sec value for V̇O_2_ during the maximal test was defined as V̇O_2max_. W_max_ was calculated by extrapolating the W-V̇O_2_ curve from the submaximal tests of each individual to V̇O_2max_.

Only in the ET participants, fat oxidation rates (g min^-1^) at submaximal power outputs (170, 205, and 240 W) were determined using the stoichiometric equation developed by Frayn (1983): fat oxidation = (1.67 V̇O_2_) j (1.67 V̇CO_2_), with urinary nitrogen excretion assumed to be negligible. Additionally, polynomial curve fitting was employed to assess the relationship between the relative utilization of V̇O_2_max during the graded test and fat oxidation rate. The relative utilizations of V̇O_2_ were segmented into 7.5% intervals from 30% to 75% of V̇O_2_max. Maximal fat oxidation was derived from the apex of the best-fit polynomial curves for each subject.

### Transmission electron microscopy

A small specimen (< mm^3^) of the muscle biopsies was fixed in 2.5% glutaraldehyde in 0.1 M Na-cacodylate buffer (pH 7.3) for 24 h at 5°C and then washed in Na-cacodylate buffer. The samples were then stored at 5°C until further processing as described elsewhere (Jensen et al. 2022). Briefly, the fixed specimens were post-fixed with 1% osmium tetroxide and 1.5% potassium ferrocyanide in 0.1 M Na-cacodylat buffer, rinsed in the Na-cacodylat buffer, dehydrated through graded series of alcohol, infiltrated with graded mixtures of propylene oxide and Epon, and embedded in 100% fresh Epon and polymerized at 60°C for 48 h. Two thin (60 nm) sections were cut by an ultramicrotome (Ultracut UCT ultramicrotome, Leica Microsystems, Wetzlar, Germany) interspaced by 150 µm in depth of the Epon block and contrasted by uranyl acetate and lead citrate. Sections were imaged in a pre-calibrated transmission electron microscope (EM 208, Philips, Eindhoven, The Netherlands) using a Megaview III FW camera (Olympus Soft Imaging Solutions, Münster, Germany). From each biopsy, all longitudinal oriented muscle fibres (n = 3-9) present in the two sections were photographed by a systematic, but random protocol ensuring unbiased results. Also, the systematic protocol ensures that images are obtained by both the subsarcolemmal (12 images per fibre) and myofibrillar regions (12 images per fibre).

Images were analyzed by manually outlining every mitochondrial and LD profile, as well as the membrane interaction between the two, using the software RADIUS (RADIUS Desktop 2.0, Build 14681, EMSIS GmbH, Germany) and the “interpolated polygon” function. The following measures were collected: volume percentage, profile area, profile aspect ratio (the maximum ratio between the length and width of a bounding box), profile convexity (the area relative to the area of the profile’s convex hull), and length of contact between the mitochondrial and LD profiles (contact strictly defined as physical contact between the two organelles). Examples of these measures are shown in Fig. 1.

**Fig 1.**
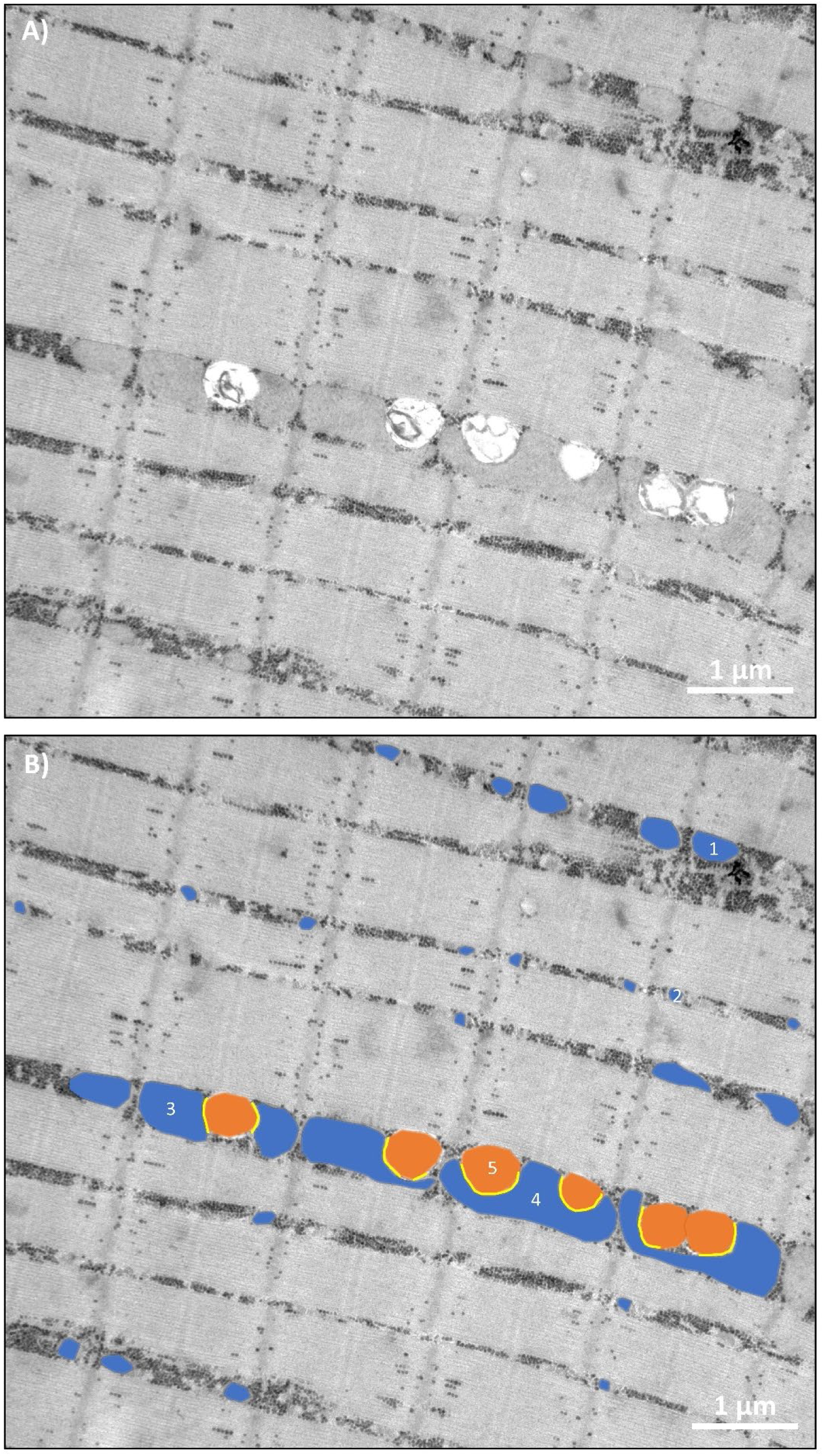
TEM images depicting the typical distribution, size, shape, and membrane interactions of mitochondrial and LD profiles. A) TEM image from an endurance-trained (ET) athlete displaying intermyofibrillar mitochondria and LDs interspersed between the myofibrils. Glycogen particles are represented by black dots. B) The same TEM image as in A), with manually outlined mitochondrial (blue) and LD (orange) profiles, along with their membrane interactions (yellow). The area (µm^2^), aspect ratio, convexity, and length of contact (nm contact for the LD) for five profiles were 1) 0.10, 2.04, and 0.99; 2) 0.02, 1.18; and 0.99; 3) 0.25, 1.76, and 0.96; 4) 0.57, 2.97, and 0.65; 5) 0.19, 1.24; 0.99, and 916, illustrating mitochondrial profiles of different sizes and shapes and a typical LD.

The differentiation between the subcellular compartments, subsarcolemma and intermyofibrillar, was performed. The volume percentages of intermyofibrillar mitochondria and LDs within a fibre were estimated by dividing the total area of the profiles by the total area of the images, following Cavalieri’s principle. This principle demonstrates that the area fraction equals the volume fraction of large structures observed in thin 2D sections. Hence, the cellular content of intermyofibrillar mitochondria and LDs is expressed as volume percentages of the myofibrillar space.

The expression of subsarcolemmal mitochondria and LDs was based on fibre surface area for two reasons. Firstly, when dealing with longitudinally oriented fibres, the fibre’s size (diameter) is unknown, which complicates estimating the contribution of the subsarcolemmal (SS) space to the overall fibre volume. Thus, expressing SS mitochondria or LD volume per fibre volume could skew the comparison of fibres, with SS contribution potentially being overestimated in thin randomly cut fibres and underestimated in thick ones. Secondly, the subsarcolemmal space contains various components such as mitochondria, LDs, glycogen, nuclei, etc., which vary greatly. Estimating a volume percentage of mitochondria or LDs per volume of the SS space would be significantly influenced by the presence of other inclusions. Consequently, the LD volume percentage might appear lower if the mitochondrial volume percentage was elevated, and vice versa. Therefore, estimating mitochondrial and LD volume per surface area remains unaffected and thus provides a robust measure.

The total fibre volume percentage of intermyofibrillar and subsarcolemmal mitochondria and LDs was estimated by converting the subsarcolemmal (SS) volume per surface area to a volume per myofibrillar volume. In this calculation, fibres were assumed to have a diameter of 80 µm and to be cylindrical in shape. Thus, one square micron of surface area corresponds to 20 µm^3^ of myofibrillar volume. Consequently, the SS volume per surface area was divided by 20 and then added to the intermyofibrillar volume percentage to obtain the total fibre volume percentage.

The average of at least seven LDs are required to obtain a reliable estimate (de Almeida et al. 2023a). This could not be met for IMF LDs in one ET participant and for SS LDs in two ET participants. Thus, the sample size for LD size, shape, and numerical density were accordingly lower.

### Confocal microscopy

Cryosections (5 µm) were cut at -30 °C and mounted on to ethanol-cleaned glass slides, fixed in 3.7% formaldehyde and permeabilized in 0.5% Triton-X 100 for 5 mins. After washing in phosphate-buffered saline (PBS) the muscle sections were incubated for 1 h with the following combination of primary antibodies: guinea pig polyclonal anti-OXPAT (PLIN5; cat no. GP31, Progen Biotechnik, Germany), and mouse anti-myosin antibody for slow twitch fibres (A4.840-c, DSHB, developed by Dr. Blau). After further washing with PBS, sections were incubated with appropriate Alexa Fluor secondary antibodies (Invitrogen, Paisley, UK) for 30 min, then washed again, followed by incubation for 5 min with BODIPY 493/503 (Invitrogen, Paisley, UK) to visualise IMTG. Vectashield mounting medium (H-1000, Vector Laboratories, Burlingame, CA, USA) was used to mount the coverslips which were subsequently sealed with nail varnish.

Images of cross-sectionally orientated sections were used to assess LD co-localisation with PLIN5 and were captured using an inverted confocal microscope (Zeiss LSM710, Carl Zeiss AG, Oberkochen, Germany) with a 40x 0.75 NA oil immersion objective and 8x digital zoom. An argon laser was used to excite BODIPY, and a helium-neon laser was used to excite the Alexa Fluor 546 and 633 fluorophores. Fibres stained positively for myosin heavy chain type I were classified as type I fibres, whereas those with no staining were classified as type II fibres; this classification was used to assess fibre-specific volume percentage of PLIN5, as previously described (Shepherd et al., 2013) (Fig. 10).

Image processing and co-localisation analysis was undertaken using Image-Pro Plus 5.1 software (Media Cybernetics, MD, USA). This method has been described in detail previously (Shepherd et al., 2013), but briefly a positive signal for PLIN5 and IMTG in sequential images was first obtained by selecting a uniform intensity threshold. Based on the selected threshold, binary images were created for PLIN5 and IMTG and the images were merged to generate a co-localisation map (Fig. 10E). The overlapping regions were subsequently extracted into a separate image (Fig. 10F), and the total number of extracted objects in this image as a proportion of the total number of PLIN5 objects was used as a measure of co-localisation. The number of extracted objects was expressed relative to area and therefore represents the density of PLIN5-associated LDs (PLIN5+ LDs). In addition, the number of extracted objects was subtracted from the total number of LDs, and expressed relative to area, to quantify the density of LDs not associated with PLIN5 (PLIN5-LDs). A total of 278 fibres were used for colocalization analysis (166 type I fibres, 112 type II fibres).

### Statistics

All effect sizes with 95% CI (Fig. 2, 5, 6 and 7) were estimated by bootstrapping (5000) using the Durga-package in R (Kahn & McLean, 2023). All scatter plots with correlation coefficients (Fig. 3, 4, 9 and 10) were conducted in Microsoft Excel^®^ and with confidence intervals of the coefficients computed as proposed by Riffenburgh (2012). Histograms (Fig. 5 and 7) were performed in Microsoft Excel. The interactions between profile areas and aspect ratio and convexity were investigated by a mixed effect model using Stata 18 (StataCorp, College Station, TX, USA) with profile area and aspect ratio or convexity as fixed effects and participant and muscle fibre as random effects. The random effects were included to take into account the nested design with multiple profiles originating from the same muscle fibres. Linear regression analyses of total contact length and numerical density of LDs (Fig. 8) were investigated by a fixed-effect regression model using Stata 18. Heteroscedasticity and normal distribution of model residuals were tested by Q-Q plots and by plotting the predicted values against the residuals, respectively. The convexity values were logit transformed before analysis. An interaction with a P-value below 0.05 was considered statistically significant. Effect sizes and correlation coefficients with 95% confidence intervals were interpreted without the use of P-values.

**Fig. 2.**
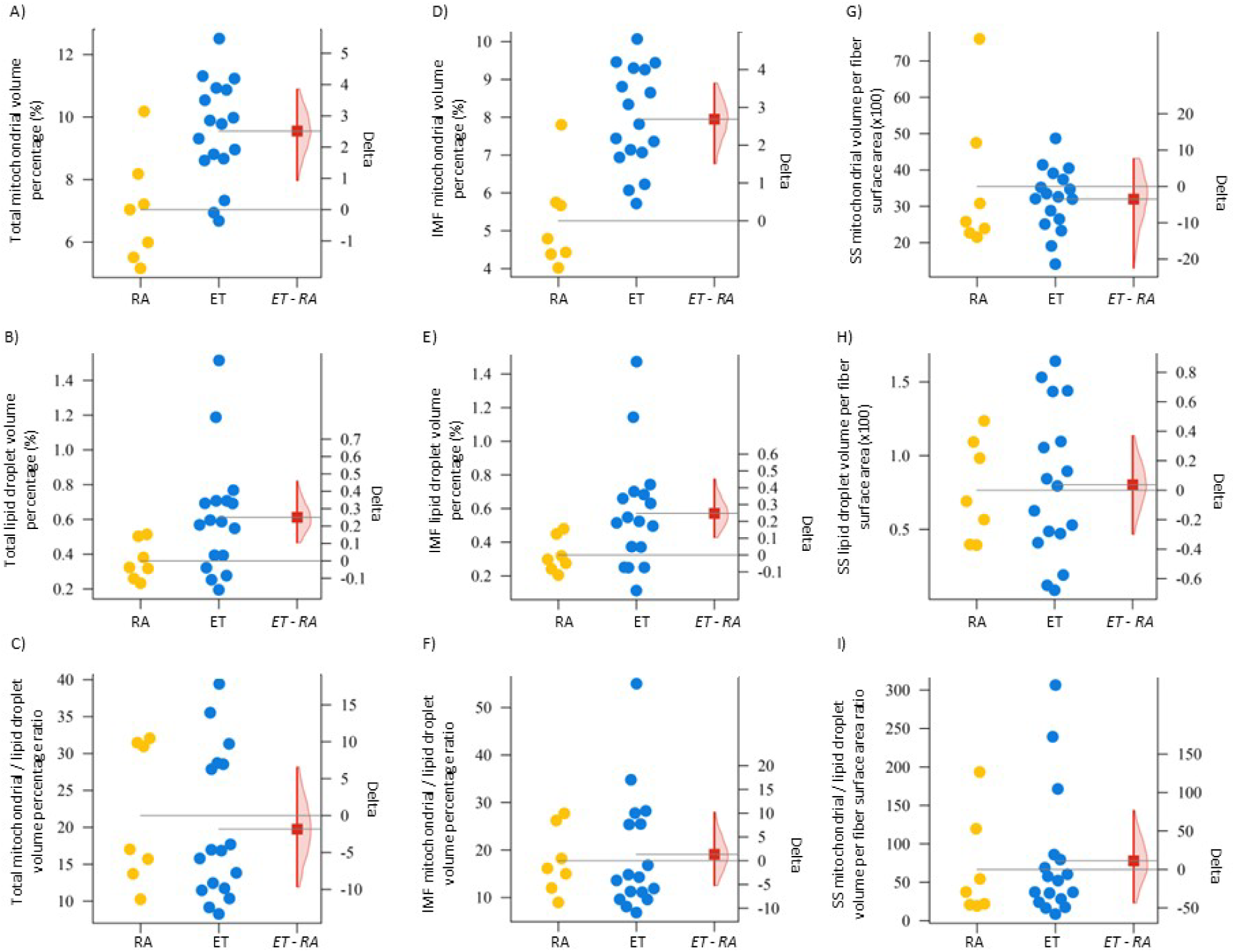
Mitochondrial and LD content in skeletal muscles of untrained and endurance-trained athletes. A-C) Total mitochondrial and LD volume percentages. D-F) IMF mitochondrial and LD volume percentages. G-I) SS mitochondrial and LD volume per fibre surface area. Each point represents one participant. RE, recreationally active (n = 7). ET, endurance-trained (n = 17). The point-estimate and 95% CI for the ET minus RA delta values were computed by bootstrapping.

## Results

### Volumetric LD and mitochondrial densities

The overall muscle fibre volume percentages of both mitochondria and LDs were markedly higher (+50%) in the ET group compared to the RA group (Fig. 2A-B). A distinction between IMF and SS subcellular localizations revealed that this difference was confined to the IMF space (Fig. 2D,G,E,H). The ratio between mitochondrial and LD volume fractions was around 20 for the IMF space and 50 for the SS space in both populations, but with high variability between individuals (Fig. 2C, F, I). Within the ET group, citrate synthase activity, but not or less clearly HAD activity, correlated with the total mitochondrial volume percentage (Fig. 3).

**Fig. 3.**
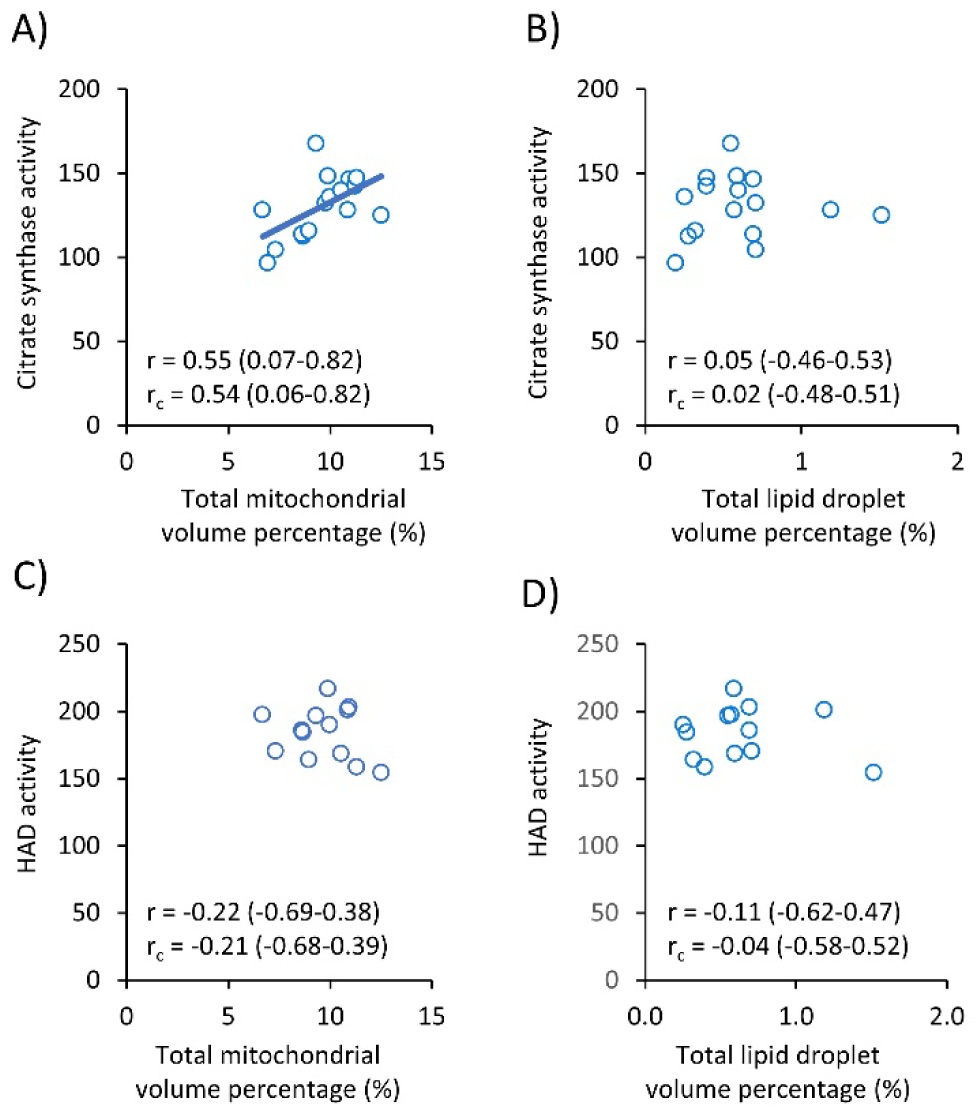
Correlations between citrate synthase and HAD activities with total mitochondrial and LD volume percentages in endurance-trained participants. A) Citrate synthase vs total mitochondrial volume percentage (n = 16). B) Citrate synthase vs total LD volume percentage (n = 16). C) HAD vs total mitochondrial volume percentage (n = 13). D) HAD vs total LD volume percentage (n = 13). Correlation coefficient (r) and concordance coefficient (r_c_) are shown with 95% CI.

Our thorough imaging of individual muscle fibres allows us to observe fibre-to-fibre variability. In the IMF space, ET athletes exhibited consistently higher mitochondrial volume percentages across the examined fibres (Fig. 4A), whereas the difference between groups in the SS space seemed most evident in fibres with the lowest mitochondrial content (Fig. 4B). The higher IMF LD volume percentage in the ET group was particularly notable in fibres with the highest LD content (Fig. 4C), which appears to correlate with a higher mitochondrial volume percentage in those fibres (Fig. 4E). Conversely, the ET group displayed a lower SS LD volume than the RA group in fibres with the least content (Fig. 4D), and this discrepancy was not or only weakly related to mitochondrial content (Fig. 4F). These observations at the single fibre level underscore a substantial fibre-to-fibre variability, which is influenced by subcellular localization and differs for mitochondria and LDs.

**Fig. 4.**
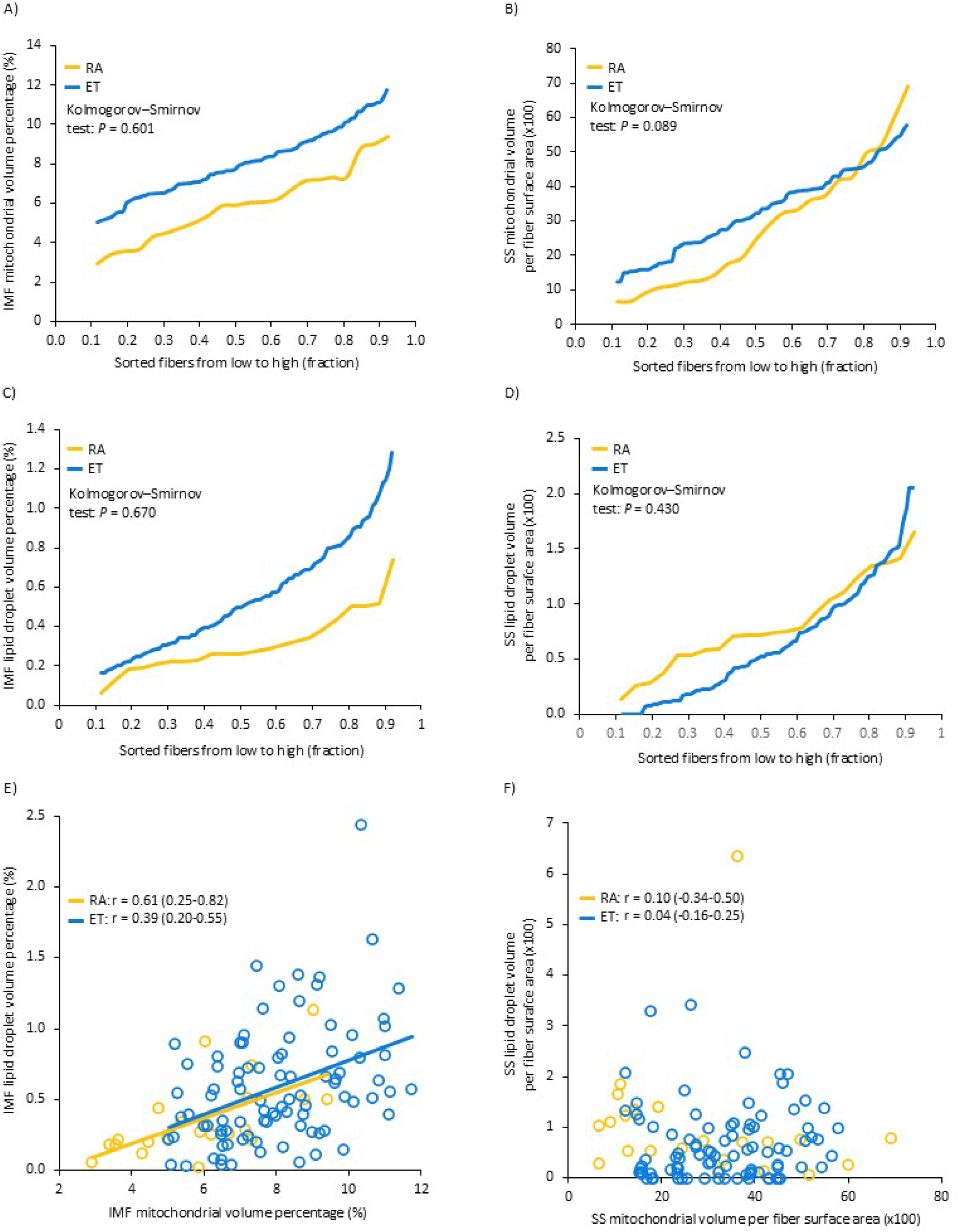
Single fibre values of IMF and SS mitochondrial and LD content. A-D) IMF and SS mitochondrial and LD content of single fibres sorted from low to high. E-F) IMF and SS LD content plotted against the mitochondrial content of the same fibre. Correlation coefficient (r) is shown with 95% CI. N = 91 and 22 single fibres from 17 and 7 ET and RA participants, respectively. In A-D, a two-sample Kolmogorov–Smirnov test for equality of distribution functions was performed using values normalised to the group mean.

### Profile size and numerical density

All mitochondrial and LD profiles were outlined manually, providing area estimates for each profile. The point estimates suggested approximately 10% larger mitochondrial profiles and 10% smaller LD profiles in the ET group compared with the RA group (Fig. 5B, E, H, K). However, considerable inter-individual variation introduces uncertainty to these point estimates, such that zero or up to 20-30% differences between the groups are also compatible with the data. The size distribution of the profiles was comparable between groups (Fig. 5C, F, I, L) and indicated a clear leftward-skewed distribution for mitochondria (Fig. 5C and 5F). In contrast, the numerical densities of IMF mitochondria and LDs were approximately 30% and 100% higher, respectively, in the ET group compared to the RA group (Fig. 5A, G) - a pattern not observed in the SS space (Fig. 5D, J). Consequently, the higher volume percentages of mitochondria and LDs in the IMF space of ET athletes (Fig. 2) can be attributed to the higher numerical densities (Fig. 5).

**Fig. 5.**
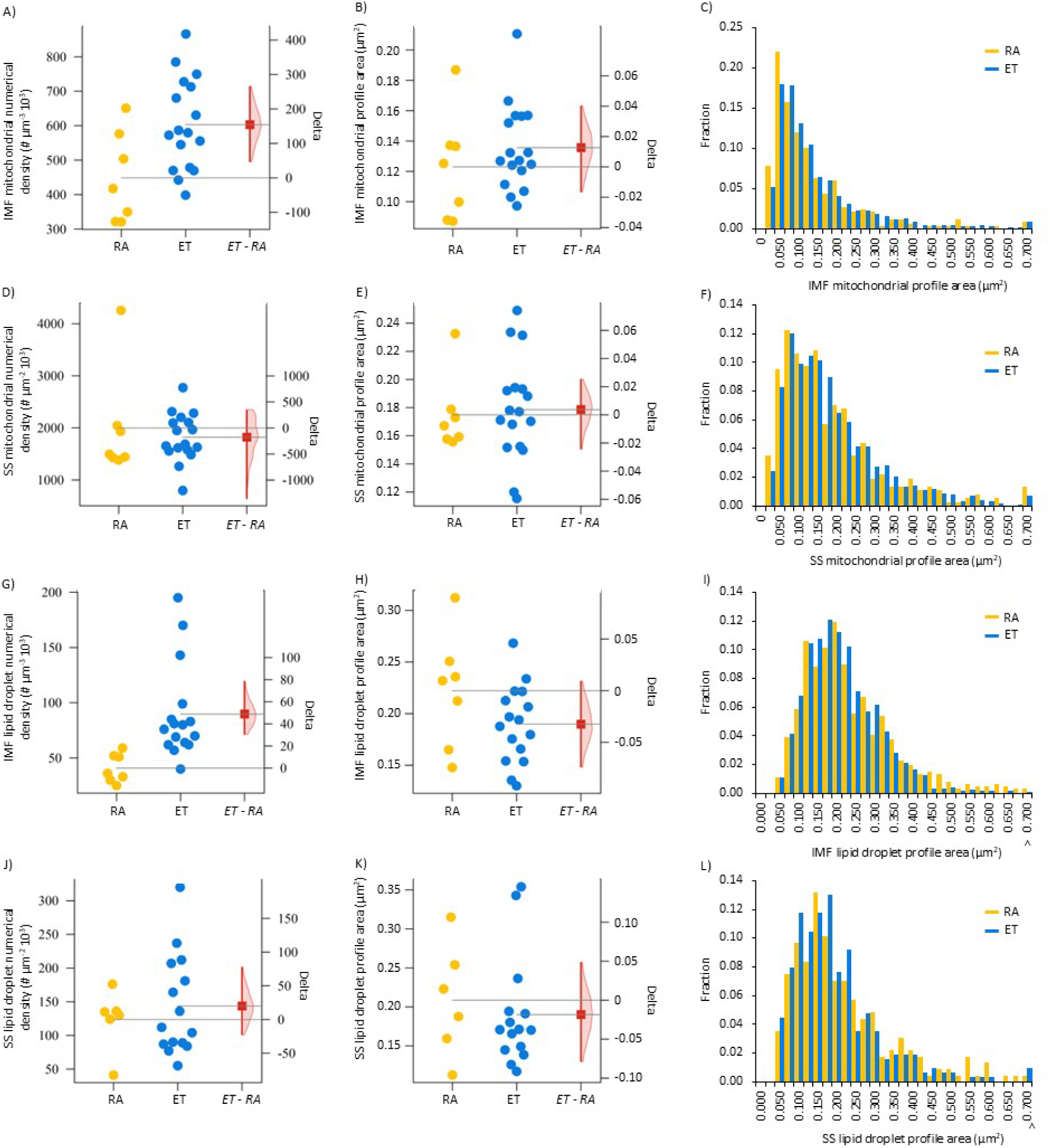
Mitochondrial and LD profile area and numerical density. A-C) IMF mitochondria. D-F) SS mitochondria. G-I) IMF LDs. J-L) SS LDs. Each point represents one participant. RA, recreationally active (n = 7). ET, endurance-trained (n = 15-17). The point-estimate and 95% CI for the ET minus UT delta values were computed by bootstrapping. Histograms of IMF mitochondrial profile areas are based on 368 and 1168 profiles from the RA and ET participants, respectively. Corresponding values were 368 and 1139, 611 and 1625, and 227 and 315 profiles for SS mitochondria, IMF LDs and SS LDs, respectively.

### Profile aspect ratio and convexity

While the aspect ratio (i.e., deviation from a circular shape) of IMF mitochondrial profiles did not differ between the groups (Fig. 6A), the SS mitochondrial profiles exhibited a lower aspect ratio in the ET group compared to the RA group (Fig. 6B). Across both groups and in both IMF and SS mitochondrial profiles, the largest profiles had the highest aspect ratio (Fig. 6A-B), indicating that the largest profiles were more frequently deviant from circular. In contrast, for LDs, the ET group showed higher aspect ratios for both IMF and SS LD profiles, with the largest profiles exhibiting the lowest aspect ratios (Fig. 6C-D).

**Fig. 6.**
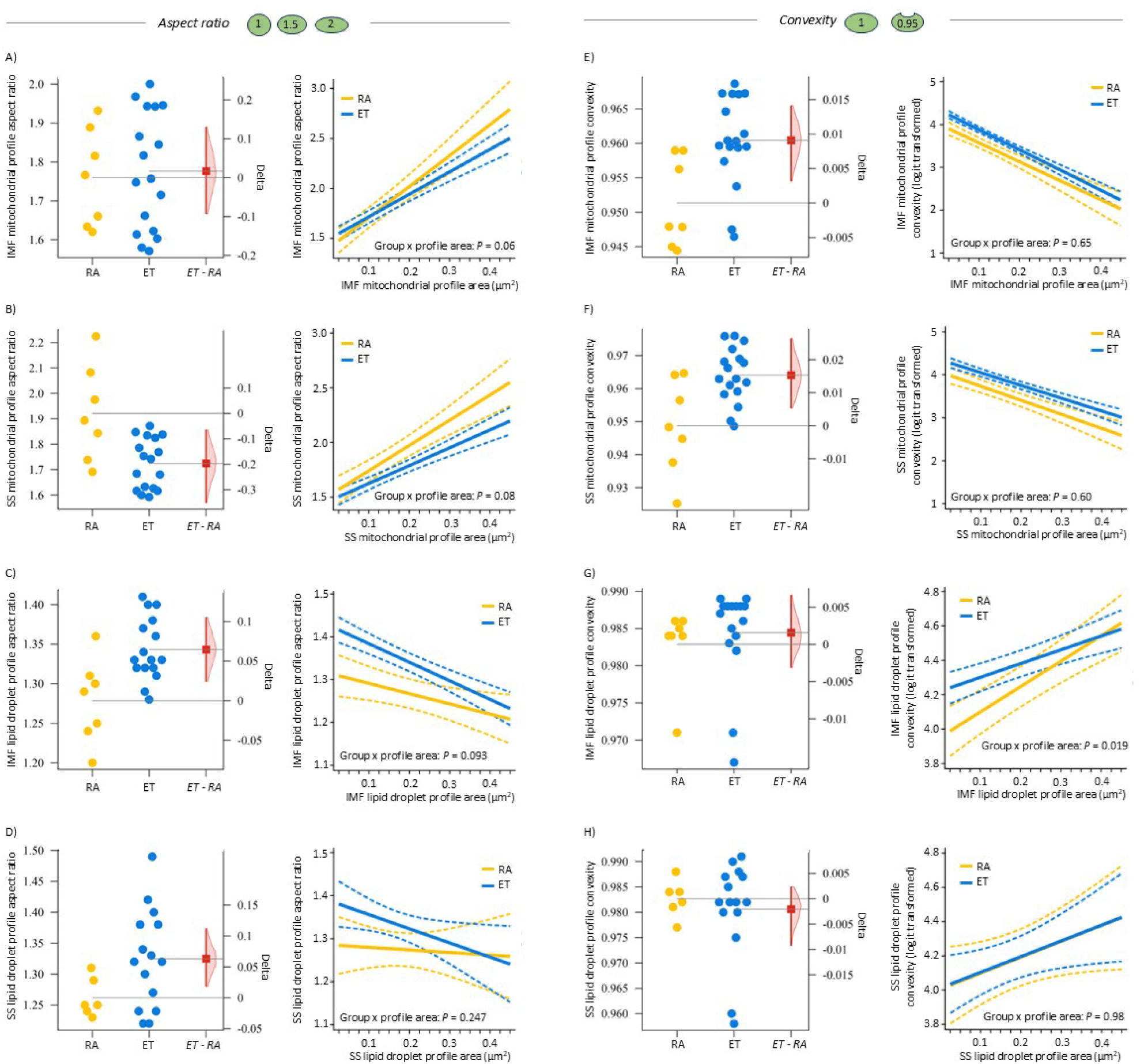
Aspect ratio and convexity of mitochondrial and LD profiles. A-D) Aspect ratio. E-H) Convexity. Each point represents one participant. RA, recreationally active (n = 7). ET, endurance-trained (n = 15-17). The point-estimate and 95% CI for the ET minus RA delta values were computed by bootstrapping. Since the size of a profiles is a strong influencer of the shape of the profile, the interaction between groups (RA vs ET) and the profile size was investigated by a mixed model including groups and profile size as fixed effects and participant and fibre as random effects. The analysis was restricted to profile areas between 0.025 and 0.45 µm^2^. The mean (and 95% CI as dotted lines) margins are plotted against the range of profile sizes. The convexity of profiles was logit transformed before the analysis and the logit transformed values are shown. N = 328 and 1068 IMF mitochondrial profiles for the RA and ET participants, respectively. Corresponding values were 336 and 1060, 575 and 1595, and 211 and 305 profiles for SS mitochondria, IMF LDs and SS LDs, respectively.

Overall, both mitochondria and LDs could generally be characterized as having a regular shape (i.e., convexity close to 1), although mitochondria profiles were often observed to be more irregular in shape compared to LDs. However, the ET group exhibited more regularly shaped mitochondria than the RA group, and across both groups, the largest mitochondrial profiles were the most irregular (Fig. 6E-F). Conversely, for LDs, the largest droplets tended to have the most regular shape. There were no large differences in the convexity of LDs between the groups (Fig. 6G-H).

### Membrane interactions between mitochondria and LDs

Approximately one-third of the LDs were not in contact with mitochondria, while of the remaining two-thirds that were in contact, the typical length of contact ranged from 150 to 750 nm. This observation held true regardless of the group or subcellular location (Fig. 7A-B, D-E). However, due to the higher numerical density of IMF LDs in the ET group, they also exhibited a 2- to 3-fold greater total contact length between LDs and mitochondria per volume of muscle (Fig. 7C, F). Since the average length of contact between individual LDs and mitochondria was not different between the two groups, the 2- to 3-fold greater total contact was explained by a higher numerical density of IMF LDs in the athletes. However, a high numerical density of IMF LDs is not unique to athletes as we have previously observed a comparable high number of IMF LDs in untrained lean individuals and obese individuals with and without type 2 diabetes (de Almeida et al. 2023). Therefore, we next included data from those additional untrained individuals in a linear regression analysis of total contact with the numerical density of LDs to investigate any potential differences in total contact length between athletes and untrained individuals with similar numerical densities of LDs. By comparing the slope of the regression lines, this revealed a 24% higher contact length in the ET than in the untrained individuals in the IMF region (Fig. 8A-B), whereas no difference was observed in the SS region (Fig. 8C-D).

**Fig. 7.**
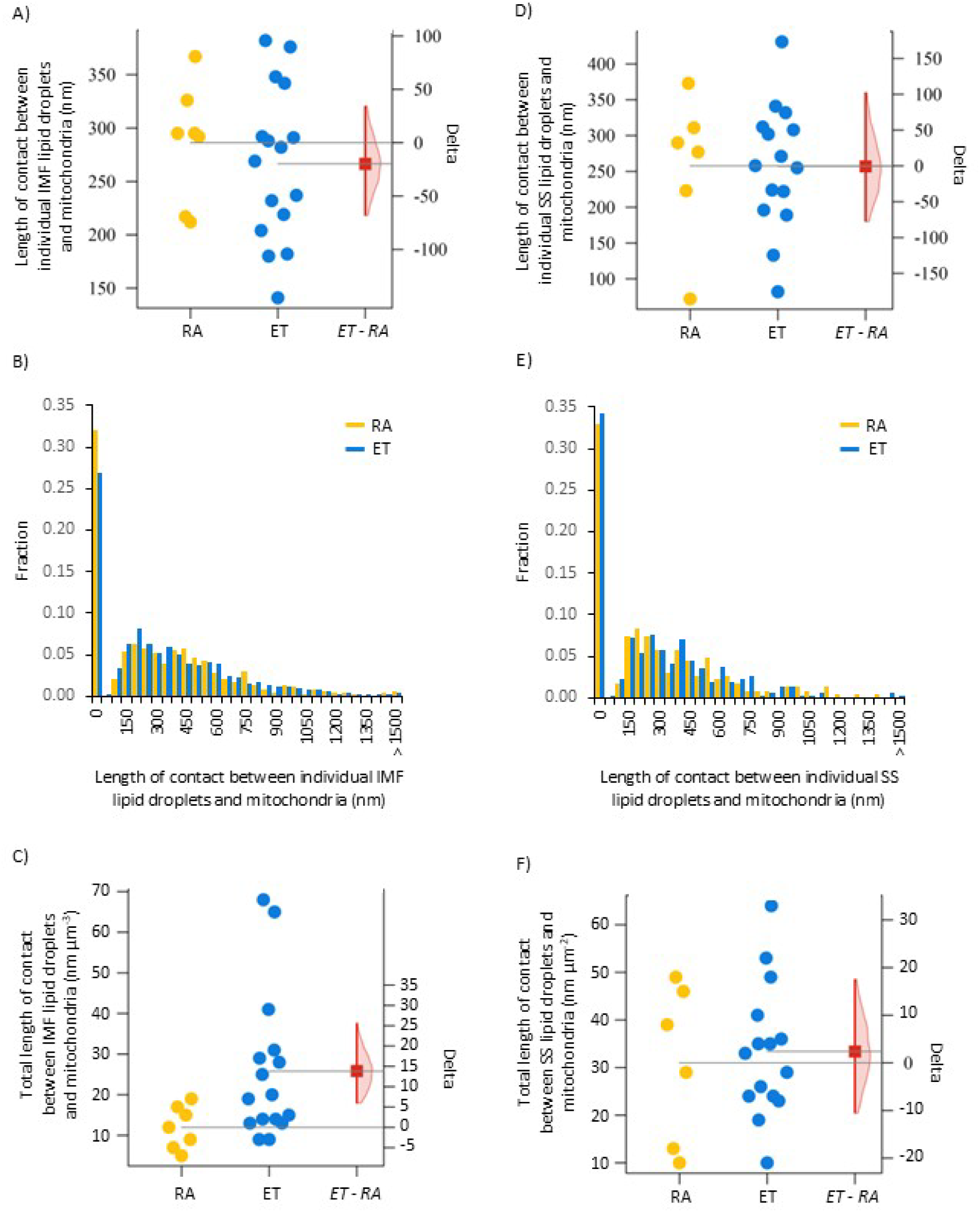
Membrane interaction between mitochondria and LDs. A-C) IMF mitochondria and LDs. D-F) SS mitochondria and LDs. In A, D, C and F, each point represents one participant. RA, recreationally active (n = 7). ET, endurance-trained (n = 15-16). The point-estimate and 95% CI for the ET minus RA delta values were computed by bootstrapping. In B (IMF) and E (SS), the number of LDs are 611 and 1625, and 227 and 315 for the RA and ET participants, respectively.

**Fig. 8.**
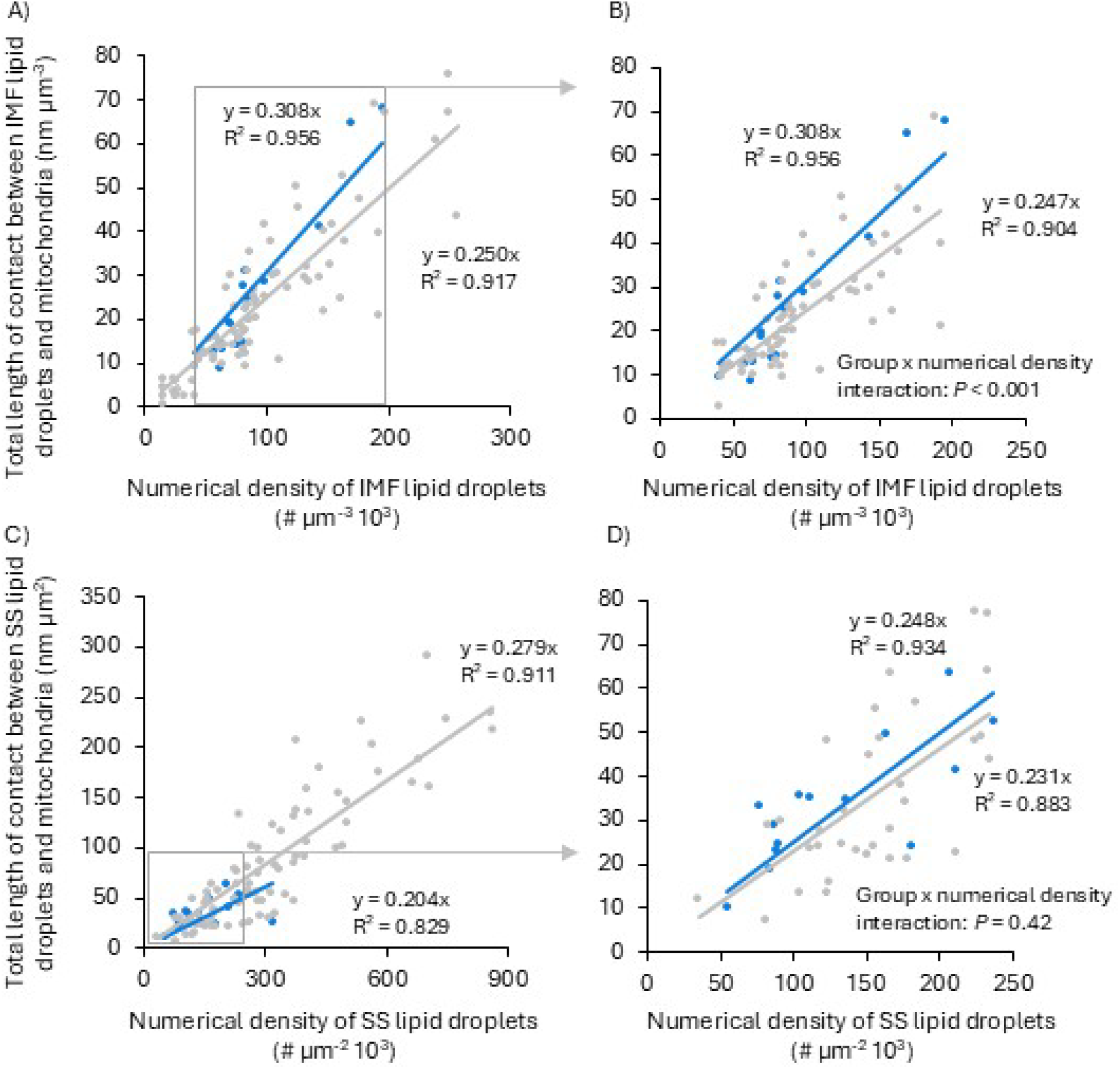
Linear regression analyses of total contact length between LDs and mitochondria with the numerical density of LDs. A-B) IMF space. C-D) SS space. In B and D) the analyses were restricted to the observations of the numerical density of LDs found in both the athletes and the untrained. The observations are grouped as the ET group (blue) and the untrained group (grey) comprising 82 observations from 43 untrained individuals from a previous cohort, where biopsies were obtained before (n = 43) and after (n = 39) 8 weeks of aerobic training (de Almeida et a. 2013a). The clustering of observations within individuals (pre and post training) was taken into account by a fixed-effect regression model. If the observations from the obese individuals with type 2 diabetes are removed from data set shown in B and D, the untrained group’s data are best fitted to y = 0.249x, R^2^ = 0.907 (two-way interaction: P = 0.002), and to y = 0.231x, R^2^ = 0.869 (two-way interaction: P = 0.408), respectively.

### Correlations with maximal fat oxidation rate and PLIN5

Within the ET group, the athletes’ maximal fat oxidation rate showed the strongest correlation with the profile size of both IMF mitochondrial and LD profiles (Fig. 9), and that a high maximal fat oxidation rate was associated with concomitant large LD and mitochondrial profiles (Fig. 9O). In the samples from the ET group, we also estimated the PLIN5 volume percentage and if LDs were PLIN5 positive or negative. The maximal fat oxidation rate showed no, or only weak, associations with both the PLIN5 volume percentage and the ratio between PLIN5-positive and -negative LDs (Fig. 10A-B). However, although the IMF LD profile area did not or only weakly correlate with the PLIN5 volume percentage (Fig. 10C), there was a moderate-to-strong correlation (if one observation was not included) with the ratio between PLIN5-positive and -negative LDs (Fig. 10D), suggesting that larger LDs are more likely to be PLIN5-positive.

**Fig. 9.**
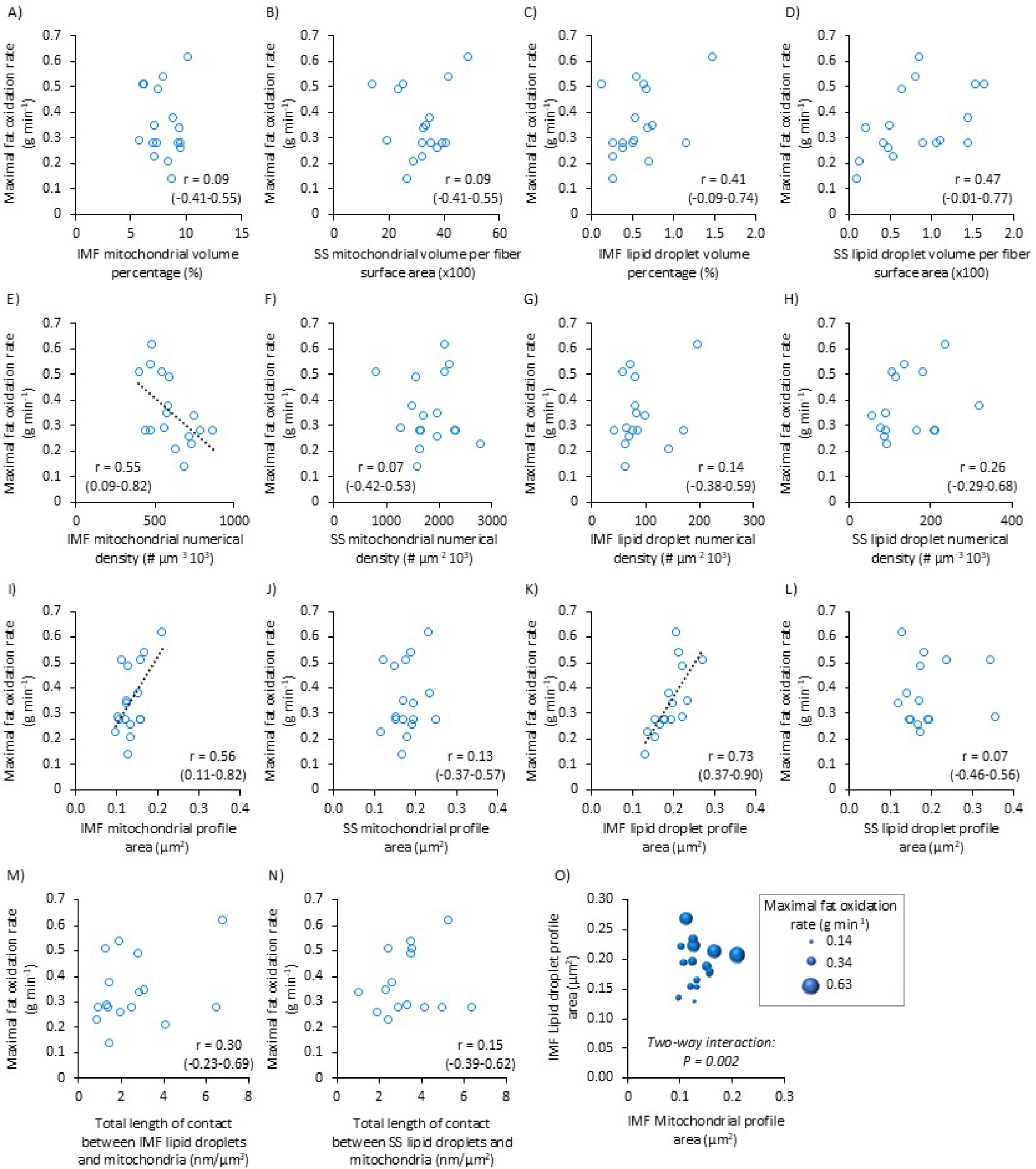
Correlations between ET participants’ maximal fat oxidation rate and mitochondrial and LD parameters. N = 17 for all mitochondrial parameters; N = 16 for IMF LD parameters; N = 15 for SS LD parameters. Correlation coefficient (r) is shown with 95% CI. If the 95% CI is above or below 0, the best fitted line is shown. In O) the maximal fat oxidation rate is shown as the size (diameter) of the spheres and with the mean IMF LD and mitochondrial profile areas shown on the y- and x-axis, respectively. The interaction was investigated by multiple linear regression.

**Fig. 10.**
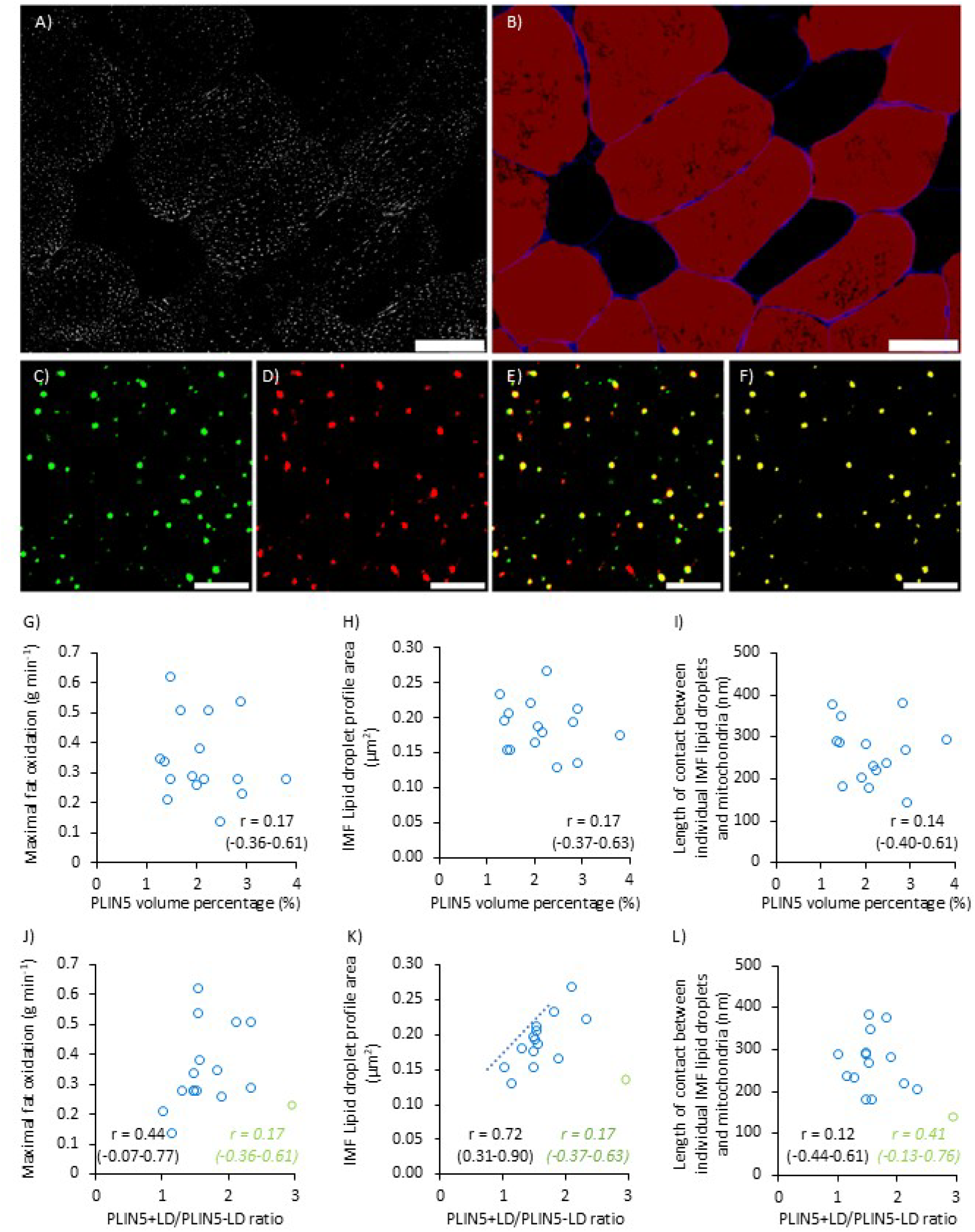
Correlations between perilipin 5 and maximal fat oxidation rate and IMF LD profile area. A-B) PLIN5 volume percentage was quantified from immunofluorescence images of PLIN5 (shown in grey scale in A), where myosin heavy chain I (MHC I) (stained red in B) was combined with wheat germ agglutinin Alex Fluor 594 (WGA) to identify the cell border (stained blue) in skeletal muscle. White bar = 50 μm. C-F) Images for colocalization analysis were obtained at 8x zoom from the central region of a cell. PLIN5 was stained in red (C), IMTG were stained with BODIPY 493/503 in green (D), and the subsequent merged images were used to calculate colocalization (E). The extracted overlying area in yellow (F) was used to calculate the relative association of PLIN5 with IMTG and to determine the number of PLIN5+ and PLIN5-LDs. White bar = 5 μm. G-L) Correlation coefficient (r) is shown with 95% CI. If the 95% CI is above or below 0, the best fitted line is shown. In J-L) r-values are shown for both all values (green) and without one outlier (blue). N = 14-16.

## Discussion

Endurance-trained athletes are characterized by a superior whole body fat oxidation capacity. Here, we show that they exhibit a greater length of contact between LDs and mitochondria than observed in recreationally active individuals and that this is confined to the intermyofibrillar space and to fibres also high in their volumetric density of mitochondria. An examination of size and shape of the organelles revealed no profound differences between the athletes and the recreationally active individuals.

## LD-mitochondria membrane contact in skeletal muscle

Our data suggest that the main determinant of total contact length between LDs and mitochondria is the numerical density of LDs and to a minor degree the contact per LD. Regarding the latter, we observed a comparable contact length per LD of around 250-300 nm in the endurance-trained and recreationally active individuals, which is greater than the 200-205 nm observed in the untrained individuals as reported in de Almeida et al. (2023a). This study is the first comparison of highly trained athletes with recreationally active individuals, but the lack of or small difference in the individual LD contact is consistent with data from the literature. Here, the most often used parameter is the percentage of LDs touching mitochondria. Using this parameter, the present results showed that the group of endurance-trained athletes had approximately 80% contact, which is comparable to or slightly higher than reported in most other studies using untrained individuals (Tarnopolsky et al. 2007; Devries et al. 2007). One study showed very low contact (25%) in untrained individuals with very low (16-20 ml O_2_ min^-1^ kg^-1^) cardiorespiratory fitness (Devries et al. 2013) and this was markedly increased after 12 weeks of endurance training, suggesting that a small dose of training is enough to increase contact (i.e., % touch). However, untrained individuals with higher fitness levels and higher contact show no large effects of short-term training on contact (Shepherd et al. 2017; Samjoo et al. 2013; Koh et al. 2018; de Almeida et al. 2023a; Devries et al. 2007). Thus, more extensive training may not influence the degree of contact.

While difficult to examine in human biopsy material, the role of contact may be to enhance the rate of fat oxidation. This is based on the finding that contact facilitates FA delivery from LDs to mitochondria (Miner et al. 2023) and that mitochondria connected to LDs have a higher respiratory capacity than LD-free mitochondria (Kim et al. 2024). In our data with the endurance-trained athletes, the total contact was not associated with the whole-body maximal fat oxidation rate. Maximal fat oxidation is dependent on several factors, and describing the minor contribution from total contact would require a much higher sample size.

## The athlete phenotype: Greater numerical density of LDs in the intermyofibrillar space of mitochondria-rich skeletal muscle fibres

A greater numerical density of LDs in the athletes compared to the recreationally active individuals is in line with the findings of Daemen et al. (2018) and aligns with the increased numerical density following endurance training or high-intensity interval training as found in most studies (Tarnopolsky et al. 2007; Samjoo et al. 2013; de Almeida et al. 2023a), but not all (Li et al. 2014; Koh et al. 2018). In contrast, one study found an increased size of individual IMF LDs following training (Devries et al. 2013). This latter study involved women exhibiting very small LDs before the training intervention, suggesting a sex-dependent regulation of the size and number of LDs.

Our findings reveal a strong relationship between the numerical density of IMF LDs and the mitochondrial density, which is also consistent with the findings of Daemen et al. (2018), who observed a higher numerical density in type 1 fibres, as well as with the correlation between IMF LDs volume and IMF mitochondrial volume in sedentary women reported by Devries et al. (2013). However, we have recently observed a comparably high IMF LD numerical density in untrained individuals, similar to that observed in the athletes of the present study (de Almeida et al. 2023a). This suggests that aerobic fitness or mitochondrial content are not the sole drivers of IMF LD expansion.

The shape of LDs was not different between the athletes and the recreationally active individuals. This aligns with the absence of a large effect of high-intensity interval training on the LD shape (de Almeida et al. 2023a). However, after acute exercise (with no net utilization of LDs) the LDs become more spherical (de Almeida et al. 2023b) and in the present investigation we observe that the larger the droplet the more spherical and regular it appears. Theoretically, larger droplets become more spherical due to a higher tension, but they also become more accessible for protein targeting (Thiam et al. 2017). We found a positive association between large droplets and the relative distribution of LDs with PLIN5, corroborating the idea that larger droplets have more binding sites for proteins (Wolins et al. 2006). Also, we found that a high maximal fat oxidation rate was associated with skeletal muscles containing large lipid droplets and large mitochondrial profiles. This is in line with the notion that PLIN5 positive lipid droplets may be preferentially utilized during exercise (Whytock et al. 2018; Shepherd et al. 2013).

The larger IMF mitochondrial volume density in the athletes compared to the recreationally active individuals was characterized by only a trivial-to-small increase in the size of the mitochondrial profiles, but a large increase in the numerical density of mitochondrial profiles. The latter indicates an increased network with a more branched mitochondrial network. This contrasts with the findings after training studies, where the size of the mitochondrial profiles has been shown to be increased, while the numerical density seems unchanged (Tarnopolsky et al. 2007; Samjoo et al. 2013; Meinild-Lundby et al. 2018). However, the numerical density seems to increase most in individuals also demonstrating a large increase in the mitochondrial volume density (Meinild-Lundby et al. 2018), suggesting that the modest increase from low to moderate volume density may be achieved by a hypertrophy of the network (increased profile size), whereas further increments are associated with a more branched network (increased numerical density in 2D microscopy images). A more branched network may be beneficial for energy transduction and local energy homeostasis (Willingham et al. 2021).

## Conclusion

Morphometric analyses of transmission electron microscopy images of longitudinally oriented skeletal muscle fibres revealed a greater contact length between LDs and mitochondria in endurance-trained male athletes compared to recreationally active men. This higher contact length was confined to the intermyofibrillar space and to muscle fibres rich in mitochondria.

## Acknowledgement

We acknowledge the Core Facility for Integrated Microscopy, Faculty of Health and Medical Sciences, University of Copenhagen.

## Author contributions

J.N. and K.D.G conceived and designed research; K.D.G., M.E.d.A, B.C., M.H.P, K.H., and N.Ø. performed experiments; J.N., K.G.P., S.O.S., and K.D.G. analyzed data; J.N., K.G.P., S.O.S., N.Ø., and K.D.G. interpreted results of experiments; J.N. and K.D.G prepared figures and wrote manuscript; J.N. and K.D.G. drafted manuscript; J.N., K.G.P., M.E.d.A., S.O.S., B.C., M.H.P., K.H., N.Ø., and K.D.G. edited and revised manuscript; J.N., K.G.P., M.E.d.A., S.O.S., B.C., M.H.P., K.H., N.Ø., and K.D.G. approved final version of manuscript.

